# High-throughput deep learning variant effect prediction with Sequence UNET

**DOI:** 10.1101/2022.05.23.493038

**Authors:** Alistair S. Dunham, Pedro Beltrao, Mohammed AlQuraishi

## Abstract

Understanding the consequences of protein coding mutations is important for many applications in biology and medicine. The vast number of possible mutations across species makes comprehensive experimental characterisation impossible, even with recent high-throughput techniques, which means computationally predicting the consequences of variation is essential for many analyses. Previous variant effect prediction (VEP) tools, generally based on evolutionary conservation and protein structure, are often computationally intensive, making them difficult to scale and limiting potential applications. Recent developments in deep learning techniques, including protein language models, and biological data scale have led to a new generation of predictors. These models have improved prediction performance but are still often intensive to run because of slow training steps, hardware requirements and large model sizes. In this work we introduce a new highly scalable deep learning architecture, Sequence UNET, that classifies and predicts variant frequency directly from protein sequence. This model learns to build representations of protein sequence features at a range of scales using a fully convolutional U-shaped compression/expansion architecture. We show that it can generalise to pathogenicity prediction, achieving comparable performance on ClinVar to methods including EVE and ESM-1b at greatly reduced computational cost. We further demonstrate its scalability by analysing the consequences of 8.3 billion variants in 904,134 proteins detected in a large-scale proteomics analysis, showing a link between conservation and protein abundance. Sequence UNET can be run on modest hardware through an easy to use Python package.

## Introduction

Proteins are integral to biology, driving all the molecular and cellular processes that create life as we know it. The key to their success is the ability to create complex properties from limited amino acid types, which allows the many and varied processes required in living organisms to be heritably encoded. Thus, understanding the impact of genotypic changes on proteins and the phenotypes they create is a major question in biology and medicine. The number of potential coding mutations in even a single protein meant it was impossible to measure all their consequences until recent multiplexed deep mutational scanning (DMS) assays^1^, and the number of genes and species means it is still impractical to measure consequences for all of them. This makes it very important to be able to predict variant effects, both to prioritise variants for experiments and for direct use in analyses.

Most traditional variant effect predictors (VEP) are based on sequence conservation, known features such as binding sites and structural models. Many prediction tools take advantage of the natural experiment performed by evolution, using multiple sequence alignments (MSAs) to measure positional variation across species or individuals and estimate variant effects. For instance, SIFT4G^2^, EVCouplings^3^ and MutationAssessor^4^ are based entirely on conservation. Structure is also thought to be an important feature because structure determines protein function. Structure based models include FoldX^5^ and RoseTTA^6^, which use force field models to estimate variants’ impact on structural stability. These have previously been limited to proteins with high quality experimental or homology models, but recent developments in structure prediction, particularly AlphaFold2^7^, makes them more widely applicable^8^. Conservation and structure can also be combined by machine learning models, alongside other knowledge such as active sites, PTMs and binding motifs. Machine learning predictors include PolyPhen2^9^, Envision^10^ and Condel^11^. VEPs like these have been very impactful, allowing larger analyses and prioritising and interpreting experiments.

Neural networks have been successfully applied to protein sequence tasks, including VEP. The lack of large scale labelled pathogenic variants makes directly training a deep learning VEP difficult, meaning they generally also use evolutionary conservation as a proxy for deleteriousness. For example, DeepSequence^12^ learns a deep generative model for a sequence family that aims to capture the probability of observing each mutant sequence and whose posteriors can therefore be used to predict pathogenicity. It was consistently found among the top VEPs across a diverse set of DMS results^12,13^. It has been refined in the Bayesian Variational Autoencoder (VAE) EVE model^14^, which outperformed previous methods on human DMS data and ClinVar variants. However, these models still require MSA based training for each protein of interest, making them very computationally demanding. Protein language models, for example UniRep^15^, AminoBert^16^ or ESM-1b^17^, also use unsupervised learning to capture position specific representations, which relate to a range of properties, including conservation, structural stability and secondary structure. They are trained to predict the identity of masked amino acids across many different sequences, meaning they learn general protein sequence properties. These produce sequence representations in a single forward pass but are still computationally intensive and often require top models to be trained for downstream applications, which is time consuming due to the large model size.

An intermediate approach between capturing variation in a single protein family, as EVE does, and a general protein language model is to predict per position variant frequencies for any sequence, using labelled MSA training data. This frequency defines the position’s position specific scoring matrix (PSSM), summarising the cross-species diversity and conservation of the sequence. This approach balances capturing additional information contained in specific MSAs with general applicability, being able to predict rapidly from any input sequence. The link with conservation means such predictions could be used to predict deleteriousness directly or the model can be further fine-tuned using smaller scale labelled pathogenicity data. We apply this approach, presenting a fast, scalable deep learning predictor, Sequence UNET, and a corresponding python package. It uses a fully convolutional architecture to predict protein PSSMs from sequence with optional structural input. The model is trained to directly predict variant frequency or to classify low frequency variants, as a proxy for deleteriousness, and then fine-tuned for pathogenicity prediction. It outperforms previous de-novo PSSM predictors, such as SPBuild, accurately classifies low frequency variants and achieves high VEP performance but with greater scalability. Further, our model has comparable performance on these tasks to models based on the much larger ESM-1b protein language model. These language models are much slower and require significantly more compute power as well as additional top model training time. We demonstrate the benefits of performance and scalability by rapidly calculating all possible variants for 904,134 proteins detected in a pan-genome proteomics analysis^18^, something that would be impossible or prohibitively time consuming with previous VEPs.

## Results

### Sequence UNET model architecture

We have developed a highly scalable VEP, Sequence UNET (**Fig. 1A**), that uses a fully convolutional neural network (CNN) architecture to achieve computational efficiency and independence from length. Convolutional kernels also naturally integrate information from nearby amino acids (**Fig. 1B**). Since long range interactions frequently generate protein properties, we also designed the model to integrate distant information using a U-shaped compression/expansion architecture inspired by the U-NET image segmentation network^19^. Max pooling creates successively smaller layers that draw information from wide regions and the final classification is built up by processing features from each depth in turn, integrating information from a wide receptive field (**Fig. 1C**). Since protein structure contains information that is extremely difficult to extract from sequence alone the network supports an optional graph convolutional neural network^20^ (GraphCNN) module to summarise positional structural features (**Fig. 1D**), which are then concatenated with the sequence input. We provide a more detailed description of the model and its inputs and outputs in the **Supplementary Information**.

**Figure 1.**
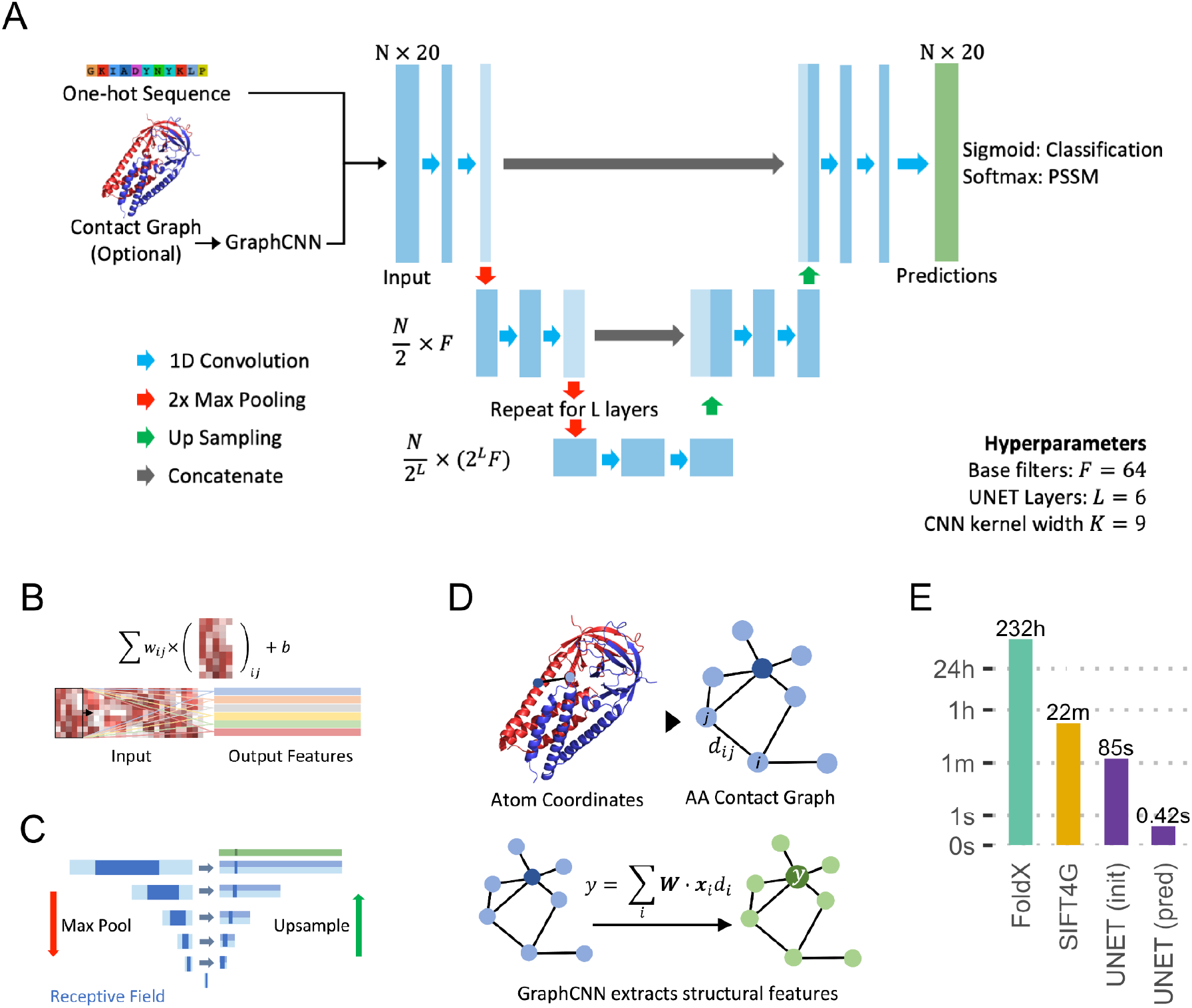
Model Overview **A**: Sequence UNET model schematic. Blue rectangles represent intermediate layer output matrices and green the final prediction. **B**: Schematic of the 1D convolution operation, which processes features in surrounding positions. Many filters (coloured arrows) are learnt in each layer to build up output features. **C**: Illustration of the receptive field in each layer from a single output position. **D**: Schematic of the optional structural GraphCNN layer. **E**: Barchart showing the computation time taken to compute predictions for all variants in SARS-CoV-2 Spike protein by two commonly used tools, SIFT4G and FoldX, and Sequence UNET. These tools were chosen for a previous analysis, but broadly span the typical timescales of current tools.

The model outputs a matrix of per position features and can therefore be trained to predict various positional properties. We demonstrate two related VEP use cases: predicting rare variants, as a proxy for deleteriousness; and directly predicting every possible variant’s frequency. We trained using ProteinNet^21^, which is a large collection of protein sequence and structure information, containing data from 104,059 structures from the PDB alongside matching variant frequencies from large MSAs based on 332,283,871 sequences. It is designed for machine learning applications and includes in-built training/validation/testing data splits based on sequence similarity and the CASP competition^22^. However, the focus on proteins with structural information may also create a bias that reduces performance on protein types that are difficult to characterise structurally. The test set is drawn from CASP12 target proteins, which have few if any related sequences included in the training data, meaning it creates a challenging test of the model’s ability to generalise to unseen and often unusual sequence space. Training is very consistent, with a variance less than 10^−5^ in both validation loss and accuracy over 10 replicates.

The trained models are highly efficient, allowing faster and larger scale prediction than comparable tools (**Fig. 1E**). For example, SIFT4G took 22 minutes and FoldX 232 hours to predict scores for all 24,187 possible variants of SARS-CoV-2 Spike protein^23^. The majority of VEPs fall within this range, with most requiring a computation intensive step such as structural sampling or MSA generation. The two most accurate neural network VEPs, EVE and DeepSequence, require both an MSA and training their latent variable model for each protein. In contrast, Sequence UNET took 420 milliseconds to compute predictions after an 85 second initialisation time (only required once per session). This enables larger scale analyses on compute clusters and rapid analyses on desktop hardware, saving valuable researcher time and resources.

### PSSM prediction and frequency classification

We trained two base Sequence UNET models, optimising performance for PSSM prediction using a softmax output layer and Kullbeck-Leibler divergence loss and variant frequency classification using a sigmoid output and binary cross entropy. Hyperparameters were tuned in both modes with the same results (selected parameters in **Fig. 1A, Fig. S1**). The PSSMs predicted by the model closely resemble true results (**Fig. S2A-B**) and the frequency classifier significantly separates rare and common variants (**Fig. S3A-B**). We find our models PSSM results correlate more strongly to true values in the ProteinNet test set than SPBuild^24^ (a state of the art de novo LSTM PSSM predictor), the amino acid propensities predicted by ESM-1b and the results from a top model using ESM-1b representations trained on ProteinNet CASP12 95% thinned data (**Fig. 2A**, **Fig. S2C-D**). Interestingly ESM-1b logits correlate much better with raw frequencies than normalised PSSMs, potentially because they are trained to identify the most likely amino acid at a position, not differentiate between the lower frequencies that are important for PSSMs. Including structural features slightly increases performance (*ρ* = 0.472 vs *ρ* = 0.451). Similar results are found for frequency classification (*f* < 0.01) over the ProteinNet CASP12 test set, with Sequence UNET achieving top performance equalling a top model using ESM-1b representations (**Fig. 2B**). We only compared to one VEP (SIFT4G) as deleteriousness is related to but not equivalent to frequency classification, instead comparing more widely on other datasets. Different frequency thresholds lead to different classification performance (**Fig. S3D**), suggesting very rare or common variants are easy to classify but intermediates are more challenging. We use the *f* < 0.01 classifier and the PSSM predictor as the base for further comparisons and generalisation. *f* < 0.01 was the most challenging threshold and a common cutoff for deleteriousness, so provides a lower bound for performance in a useful context. Further details on hyperparameter tuning and base model PSSM prediction and frequency classification performance are available in the **Supplementary Information**.

**Figure 2.**
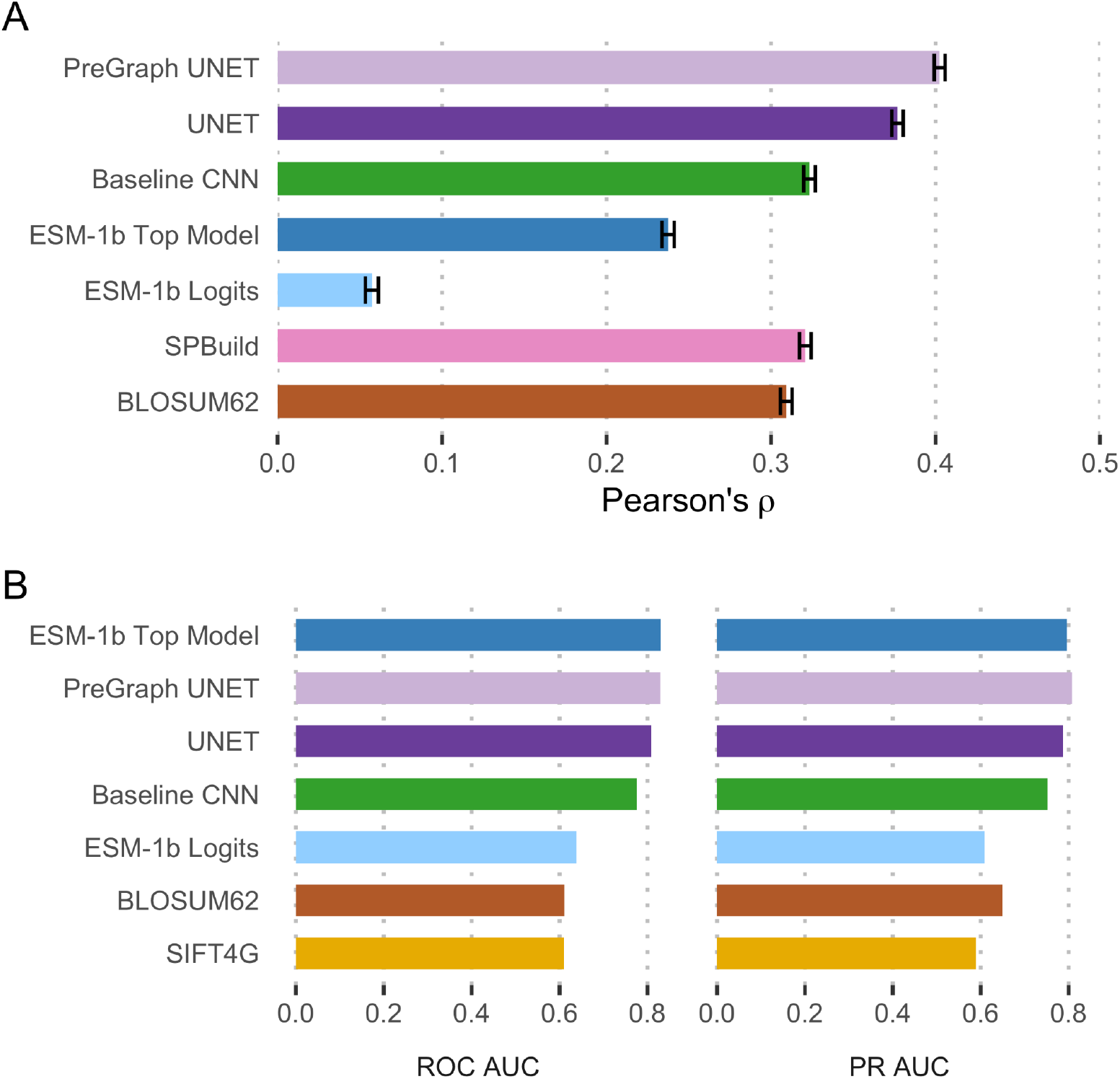
Base model prediction performance. **A**: Pearson correlation between predicted and true PSSM values comparison PSSM prediction performance for Sequence UNET, a single layer CNN, SPBuild, ESM-1b logits, an ESM-1b top model and BLOSUM62. **B**: ROC and PR curve AUC values comparing frequency classification performance of Sequence UNET with and without structural features, ESM-1b logits, an ESM-1b top model, a baseline single layer CNN, SIFT4G and BLOSUM62. All comparisons were made over the ProteinNet CASP12 test set.

### Generalising Sequence UNET

Having shown good PSSM prediction and classification performance we next sought to show Sequence UNET generalises to predicting deleterious variants and compare performance to other tools. We tested generalisation on three datasets: labelled human protein variants from ClinVar, standardised deep mutational scanning (DMS) data^25^, and a set of gold standard *S. cerevisiae* variant classifications^26^.

The model can also be fine-tuned to new tasks with additional training on external data, either refining the existing weights (fine-tuning) or replacing the final classification layer with a freshly initialised one (a top model). We trained ClinVar classification top models and fine-tuned models with and without structural features on top of the Sequence UNET frequency classification model, using a random 95%/0.5%/4.5% training/validation/testing split across all pathogenic and neutral variants in the ClinVar dataset that occur in proteins in ProteinNet CASP12 training data. The weights of all but the top two model layers were frozen to prevent overfitting. We also trained simple single layer CNN models to predict ClinVar pathogenicity and frequency classification, to provide a lower bound for machine learning solutions to this problem. The fine-tuned models specifically predict pathogenicity probabilities for each variant at all positions (**Fig. 3A**). These predictions tend to be more similar for variants at a position, including the wild-type amino acid than the results of the frequency classification and PSSM models. This is partially because pathogenicity is related to the position’s properties and importance but also suggests there might not be sufficient training data available to differentiate between different variants at one position beyond the average properties of that position.

**Figure 3.**
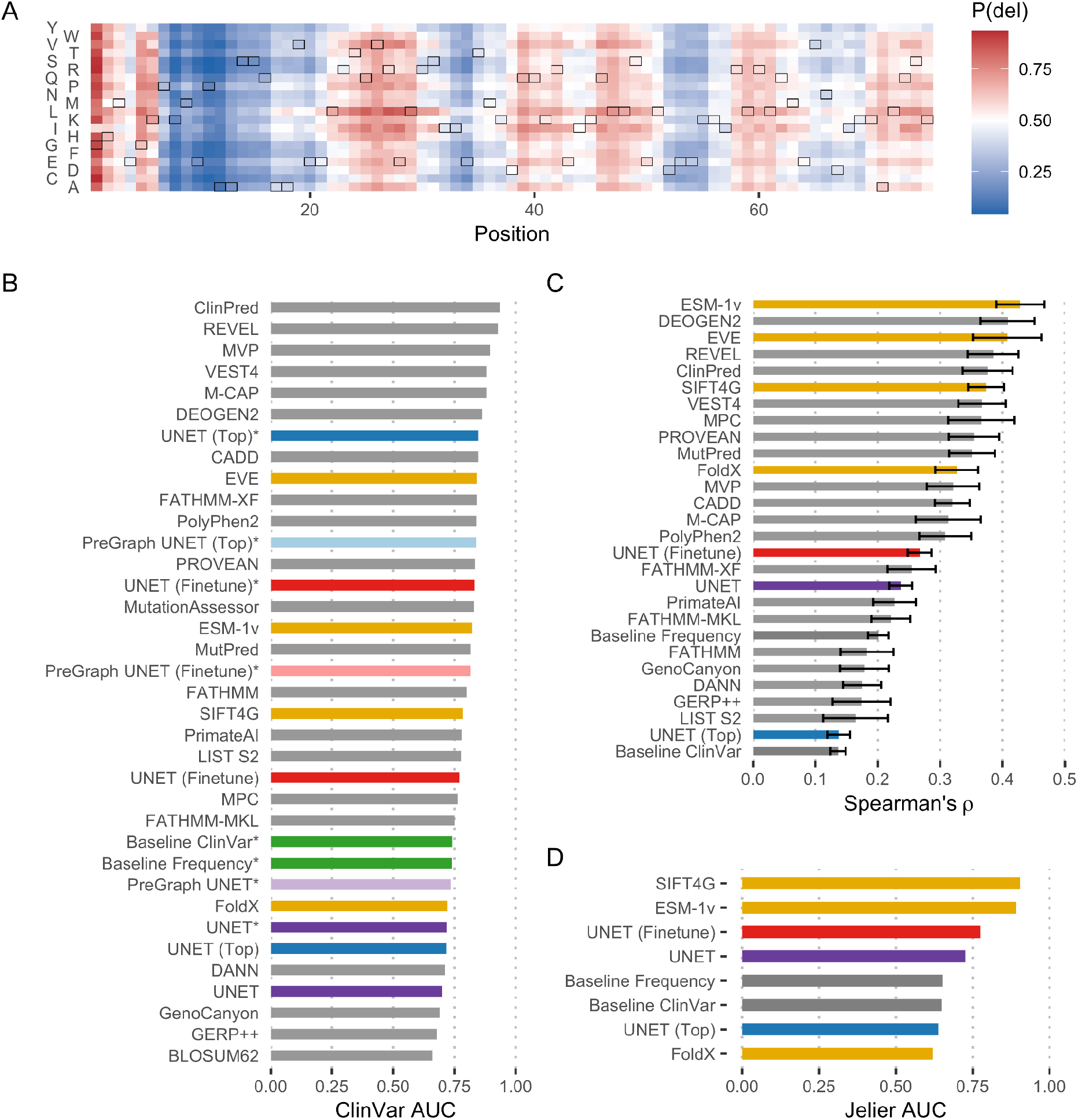
Generalising Sequence UNET **A**: Sequence UNET top model ClinVar pathogenicity predictions for the ProteinNet Casp12 test set record TBM#T0865. The wild-type amino acid at each position is outlined. **B**: ROC AUC values comparing VEP performance over the ClinVar test set. Variants of Sequence UNET are coloured in purple, blue and red, single layer CNN models in green and several notable models in yellow. Instances tested on the subset of ClinVar with structural data are marked with asterisks (*). **C:** Mean and standard error of Spearman’s rank correlation coefficient between VEP predictions and standardised DMS data^25^. Sequence UNET, ESM-1v, SIFT4G and FoldX predictions were available across all proteins while other tools were only available for human proteins. **D:** ROC AUC values comparing performance of VEPs at classifying known deleterious and neutral *S. cerevisiae* variants^26^.

Top models and fine-tuned models achieve comparable performance profiles to many state of the art predictors (**Fig. 3B**). Performance is significantly better when only proteins with structural information are considered, suggesting this bias from the ProteinNet training data has impacted what the model learnt. A training set that was not restricted to proteins from the PDB would likely help rectify this bias. The top model performs slightly better than the finetuning approach on variants with structure but worse on those without it, independently of whether the network utilised that information. This is perhaps because the freshly initialised final weights allow the network to learn new relationships specific to structured pathogenicity whereas fine-tuning maintains more general relationships. The fine-tuned UNET model only performs slightly worse than the much larger ESM-1v language model^27^ across all variants and performs better on variants with structural data, despite being a much smaller and more manageable network. Interestingly, actually utilising structural data slightly reduces the performance of the fine-tuned models, suggesting it is less related to pathogenicity. The base Sequence UNET frequency classification models generalise less well to this task, although still with comparable performance to models such as FoldX. This suggests the frequency-based model does not fully capture deleteriousness without fine-tuning.

The frequency classification model was initially trained with a range of frequency thresholds and the resulting models have different ClinVar generalisation performance (AUC_0.01_ = 0.73, AUC_0.1_ = 0.67, AUC_0.0001_ = 0.66, AUC_0.001_ = 0.65). This shows that the chosen threshold does impact performance on a given task and suggests 0.01 is the best threshold for pathogenicity prediction, which aligns with the fact that the mean allele thousand genomes frequency for benign variants is 0.112 and for pathogenic variants is 0.008^28^.

Sequence UNET predictions would also be expected to relate to DMS results and confirmed neutral and deleterious *S. cerevisiae* variants. Comparing the distribution of Spearman’s rank correlation values across DMS datasets (**Fig. 3C**) shows the model generalises, although it performs slightly less well than top predictors on this task. A similar result is found with a ROC analysis of S. *cerevisiae* variants (**Fig. 3D**), where Sequence UNET variations outperform FoldX but fall behind SIFT4G and ESM-1v. The model fine-tuned on ClinVar performs best in both cases, while the top model performs relatively poorly, even falling behind simple CNN models in some cases. This further suggests that the top model may be learning something more specific to ClinVar, which may be an artefact of the dataset but could also be a real biological feature of pathogenicity or human proteins. Together this confirms that the models generalise well to other contexts, although their relative performance compared to other tools varies.

### High-throughput proteome scale predictions

Modern high-throughput experimental approaches can generate very large quantities of data, requiring efficient computational approaches to process. For example, a recent pan-proteome analysis by Muller et al.^18^ collected protein abundance measurements from 103 species, detecting a total of 904,134 distinct proteins (**Fig. 4A**). Analysing this many proteins with the most commonly used predictors is very computationally intensive and would be prohibitively time consuming for many tools and research groups. For example, making predictions for 161,825 variants across just 30 proteins as part of a combined deep mutational scanning analysis^25^ took SIFT4G 14.1 hours and FoldX 64.5 days of total compute time. To exemplify the scalability of Sequence UNET we made predictions for all 8.3 billion possible variants in this proteomics dataset, which took 1.5 hours on a GPU using a batch size of 100 (6.8 hours without batching) and 50.9 hours using only CPU (**Fig. 4B**). The additional padding required to batch different length proteins was found to have a negligible impact on predictions for an analysis of this scale, although it does impact a small number of individual results (**Fig. S4**). This also compares favourably with the ESM language model, even when using a single forward pass for all variants instead of independently masked passes for each variant as suggested^27^. ESM is both a much larger model and the attention mechanism scales quadratically with protein length, whereas the convolutional design of Sequence UNET scales linearly. This means Sequence UNET is significantly faster using CPU and for larger proteins on GPU. Small proteins are predicted at a similar rate on GPU, suggesting at this point other factors dominate. Sequence UNET also requires much less (V)RAM in all cases, making it significantly easier to deploy at scale and allowing batches of proteins to be processed simultaneously to increase efficiency. In contrast, even a batch size of 2 was prohibitive for ESM. These performance increases, combined with the Sequence UNET python package, makes large scale analyses more accessible, especially for those without high performance compute facilities.

**Figure 4.**
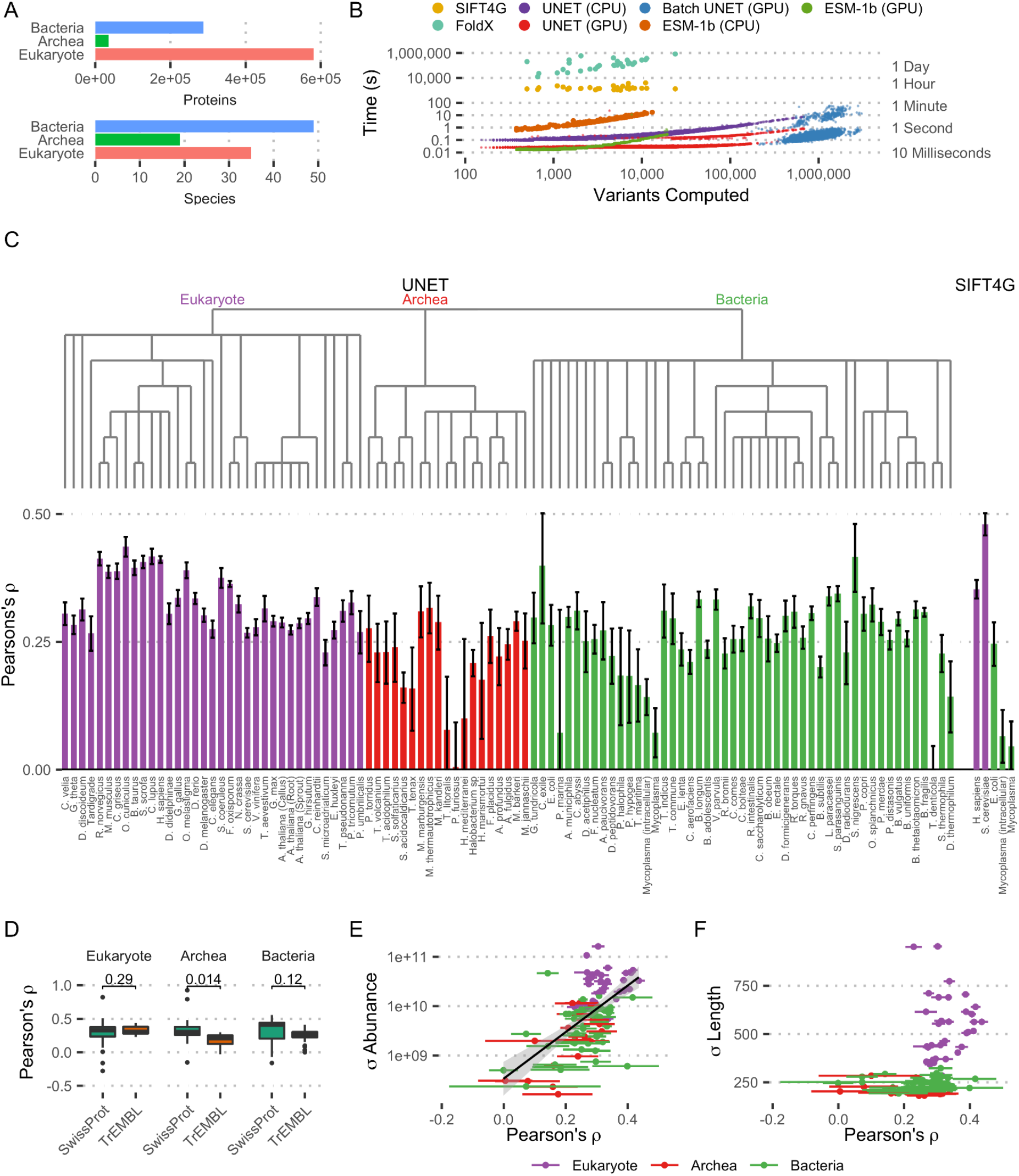
High-throughput proteome analysis **A**: Number of proteins and species in the Muller et al. proteomics dataset. **B**: Computation speed comparison between SIFT4G, FoldX, single-pass ESM-1b and Sequence UNET on CPU and GPU. Sequence UNET was also tested running on GPU in batches of 100. The SIFT4G and FoldX computations were performed as part of an independent deep mutational scanning analysis^25^, ESM-1b was run on ProteinNet proteins and Sequence UNET computations are across this proteomics dataset. **C**: Pearson correlation coefficient between predicted conservation and protein abundance in each species. The error bounds of Pearson’s ρ are calculated with Fisher’s Z transform. Predicted conservation is summarised as the mean number of variants predicted to be deleterious across positions. Results are shown for Sequence UNET frequency predictions across all species and SIFT4G for Mycoplasma and species with data available in Mutfunc^30^. The species’ phylogeny is also shown based on NCBI Taxonomy Common Tree. **D**: Boxplot showing distribution of correlation coefficients for each domain, split between proteins in SwissProt and TrEMBL. The p-value comes from a two-sample unpaired T-Test. **E**: Relationship between Pearson correlation and standard deviation of raw protein abundance across species. **F**: Relationship between Pearson correlation and standard deviation of protein length across species.

We used this large dataset of variant effect predictions for almost 1M proteins to compare protein abundance and predicted tolerance for sequence variation. Proteins that are expressed at higher abundances are generally expected to have more strongly constrained sequences than low abundance proteins^29^. This is thought to occur because highly expressed proteins need highly optimal sequences to avoid aggregation potentially driven by translation errors. However, most studies comparing abundance with sequence constraints have relied on a small number of species. In our analysis we found a significant correlation between protein abundance and predicted protein conservation in most of the 103 species in this dataset (**Fig. 4C**). Protein abundance is normalised against length and expressed as the log2 fold change compared to median abundance in that species. Conservation is summarised for a protein as the mean number of predicted deleterious variants across positions. A similar correlation level is observed for Sequence UNET predictions, including PSSM prediction and frequency classification, and SIFT4G scores in *H. sapiens*, *S. cerevisiae*, *E. coli* and Mycoplasma proteins. The similarity between correlations based on SIFT4G scores and our predictions validates the use of Sequence UNET for such applications.

The strength of conservation-abundance correlation varies a lot between species, including diminishing to nothing in a few cases. Eukaryotes tend to have the highest overall correlations (t-test vs Bacteria: *p* = 7.6 × 10^−6^, vs Archaea: *p* = 1 × 10^−5^). However, looking more closely suggests this is partly caused by the fraction of TrEMBL proteins, which may include spurious open reading frames and, in our analysis, tend to have weaker correlations in Archaea (**Fig. 4D**). The variation in protein abundance and protein length (**Fig. 4E**) also impact the abundance correlation, suggesting that part of the difference may come from reduced overall variation in protein forms. Species with no significant correlation tend to be unusual organisms, for example the intracellular parasite Mycoplasma or extremophile archaea such as *P. furiosus* or *T. litoralis*, which would be expected to have unusual properties based on their biology. They may also contain more proteins without similarities to those in our PDB based training data. There is also a stronger correlation in Mycoplasma when cultured intracellularly, suggesting it behaves more conventionally in that state and is more abnormal outside the cell. Finally, a similar correlation between abundance and conservation determined by SIFT4G scores shows a similar pattern in Mycoplasma, although with lower intracellular correlation, suggesting the low correlations have biological rather than technical causes. This simple analysis demonstrates the utility of performant, scalable predictors for large analyses and working with high-throughput experimental results.

## Discussion

Variant effect prediction (VEP) is a central part of many computational genetic analyses, allowing researchers to assess new genomes or patient sequences, prioritise variants for follow up experiments and identify important functional and structural features of proteins. This has led to a range of VEP tools, using empirical and machine learning methods to predict deleteriousness from sequence and structure. However, most tools are restrictive to run, particularly for large scale analyses involving hundreds of proteins. They can be computationally expensive, often having slow multiple sequence alignments or more recently machine learning model training as a limiting step, and are often awkward to install correctly. Precomputed results are available for human proteins and some model organisms, for example in the MutFunc^23,30^ or dbNSFP^31,32^ databases and the webpages of many tools. Despite this, it is still difficult for many scientists to generate predictions for the proteins they need, especially when analysing a large number of proteins or less studied organisms. In an attempt to fill this niche, we developed a versatile neural network VEP that is fast, lightweight and easy to use, able to rapidly make exhaustive predictions about many proteins on a regular laptop and scale to multi-proteome analyses on compute clusters.

Neural networks provide a powerful approach to fast, high-quality predictions, utilising large datasets and a long, computationally intensive training process to distil complex relationships into numerical weights matrices. This means predictions can be made efficiently as long as the network architecture uses optimised operations and doesn’t require expensive external computations, for example multiple sequence alignment or structure relaxation. Neural network models have also shown very high performance in other sequence based tasks^33–35^ and variant effect prediction^12,14^ but there has not previously been a fully end-to-end neural network specialised VEP that operates directly on sequences. The Sequence UNET model architecture we arrived on takes inspiration from other CNN models, including from sequence, structure and image based tasks^19^, and combines them into a novel model. The U-shaped compression/expansion structure allows information to propagate across the protein, with the “receptive field” of neurons in the lower layers covering large regions of the original sequence in the same way they integrate information across images in the original UNET. This allows performant CNN operations to be used for a sequence-based problem while allowing filters to learn sequence patterns at different detail levels. Similarly, GraphCNNs are performant and have been shown to perform on protein structure tasks^36–38^, and their position invariance makes them a natural approach to including structural features. We experimented with various other methods of including structure, including torsion angles and calculated feature profiles, but found GraphCNNs gave best performance and efficiency.

The lack of labelled deleterious variant data at the scale required for deep learning led us to first capture general sequence and variant properties by training the model to predict variant frequencies, either as a PSSM predictor or a low frequency variant classifier, and then finetune for pathogenic variant classification using ClinVar data. This is similar to the protein language model paradigm, in which large models are first trained to predict amino acid sequences, capturing general properties, and the representation vectors they produce can be used as input into smaller downstream models. However, the size and design of the model makes it much more computationally efficient than most language models.

The base Sequence UNET model achieves state of the art performance at de novo PSSM prediction and frequency classification, both independently useful tasks, and the fine-tuned models reach top level performance at pathogenicity prediction, although the base model only generalises moderately well. There are also potential questions about performance on unstructured proteins, which are missing from ProteinNet, but this could be addressed by expanding the training data beyond structured proteins.

Variant frequency can be measured analytically and other models can predict pathogenicity to a broadly similar accuracy, meaning the major strength of our model is computational efficiency. This enables analyses to be completed more rapidly and on weaker hardware and opens up potential large-scale analyses that would not be possible for more computationally demanding tools. For example, the multi-proteome analysis we performed here would have been extremely computationally expensive with many tools. Other possible applications include metagenome and microbiome analysis, where large numbers of new sequences are determined and need to be understood. Developing efficient models for structure, function or localisation prediction would also enhance work where a great many protein sequences are generated or need to be compared. Such methods can also be combined with slower, more accurate or analytical methods to identify the most important targets for detailed, computation intensive analysis, which gets the benefits of speed and performance by enabling accurate results from the important areas of a wide search space.

There are various routes available that could improve our model in future, while maintaining computational efficiency. The role of protein structure is an obvious target for change, since it currently only adds a small performance boost despite structure being known to be critical for protein function. For example, a richer graph network section, graph attention mechanisms or pre-training the structural section to encode structural properties could all potentially improve performance. Removing the structure option altogether could also be beneficial because it would allow the model to be trained on a much larger sequence dataset, exposing it to more sequence variation and reducing the bias towards protein types with determined structures. Large sequence datasets, and lots of training data in general, has been found to greatly improve performance in many other models^7,17,39^. This would make the model setup more similar to protein language models, which also train to predict amino acid propensity on large sequence databases. The UNET sequence CNN portion of the network could also be adjusted, either by tweaking the current connections and parameters or switching to an alternate sequence processing architecture. For instance, powerful attention and transformer architectures could be incorporated into the network or used as a basis for a new model that maintains computational efficiency as a goal, although it does come with an inherent computation cost compared to convolution.

We have demonstrated a highly efficient, performant model for variant frequency and effect prediction, which enables larger scale analyses than have previously been possible with VEP packages, demonstrated by our multi-proteome conservation analysis. This could be beneficial for a range of biological problems where current models are slow enough to be prohibitive, for example metagenomics, microbiome research and analysing the large quantities of genomic sequences from the Darwin Tree of Life project^40^. More generally, developing computationally efficient deep learning models that maintain high performance has great potential in other problems, speeding up predictions and providing approximate solutions where analytical approaches are prohibitive. This could enable new questions to be answered as well as making current analyses more economical, reducing compute time and consequently saving money and natural resources, including carbon^41^.

## Code and Data Availability

A python package and weights for easily implementing the model as well as development code are available at github.com/allydunham/sequence_unet. This code fully defines the network and training procedures. We also developed a python package for loading and manipulating ProteinNet data (ProteinNetPy), which is available at github.com/allydunham/proteinnetpy.

## Methods

Detailed methods and analysis are contained in the accompanying **Supplementary Information**.

## References

1. Fowler, D. M. & Fields, S. Deep mutational scanning: a new style of protein science. Nat. Methods 11, 801–807 (2014).

2. Vaser, R., Adusumalli, S., Leng, S. N., Sikic, M. & Ng, P. C. SIFT missense predictions for genomes. Nat. Protoc. 11, 1–9 (2015).

3. Hopf, T. A. et al. The EVcouplings Python framework for coevolutionary sequence analysis. Bioinformatics 35, 1582–1584 (2019).

4. Reva, B., Antipin, Y. & Sander, C. Determinants of protein function revealed by combinatorial entropy optimization. Genome Biol. 8, R232 (2007).

5. Schymkowitz, J. et al. The FoldX web server: an online force field. Nucleic Acids Res. 33, W382–W388 (2005).

6. Kellogg, E. H., Leaver-Fay, A. & Baker, D. Role of conformational sampling in computing mutation-induced changes in protein structure and stability. Proteins 79, 830–838 (2011).

7. Jumper, J. et al. Highly accurate protein structure prediction with AlphaFold. Nature 1–11 (2021) doi:10.1038/s41586-021-03819-2.

8. Akdel, M. et al. A structural biology community assessment of AlphaFold 2 applications. 2021.09.26.461876 https://www.biorxiv.org/content/10.1101/2021.09.26.461876v1 (2021).

9. Adzhubei, I. A. et al. A method and server for predicting damaging missense mutations. Nat. Methods 7, 248–249 (2010).

10. Gray, V. E., Hause, R. J., Luebeck, J., Shendure, J. & Fowler, D. M. Quantitative Missense Variant Effect Prediction Using Large-Scale Mutagenesis Data. Cell Syst. 6, 116–124.e3 (2018).

11. González-Pérez, A. & López-Bigas, N. Improving the Assessment of the Outcome of Nonsynonymous SNVs with a Consensus Deleteriousness Score, Condel. Am. J. Hum. Genet. 88, 440–449 (2011).

12. Riesselman, A. J., Ingraham, J. B. & Marks, D. S. Deep generative models of genetic variation capture the effects of mutations. Nat. Methods 1 (2018) doi:10.1038/s41592-018-0138-4.

13. Livesey, B. J. & Marsh, J. A. Using deep mutational scanning to benchmark variant effect predictors and identify disease mutations. Mol. Syst. Biol. 16, e9380 (2020).

14. Frazer, J. et al. Disease variant prediction with deep generative models of evolutionary data. Nature 599, 91–95 (2021).

15. Alley, E. C., Khimulya, G., Biswas, S., AlQuraishi, M. & Church, G. M. Unified rational protein engineering with sequence-only deep representation learning. bioRxiv 589333 (2019) doi:10.1101/589333.

16. Chowdhury, R. et al. Single-sequence protein structure prediction using language models from deep learning. 2021.08.02.454840 https://www.biorxiv.org/content/10.1101/2021.08.02.454840v1 (2021).

17. Rives, A. et al. Biological structure and function emerge from scaling unsupervised learning to 250 million protein sequences. Proc. Natl. Acad. Sci. 118, e2016239118 (2021).

18. Müller, J. B. et al. The proteome landscape of the kingdoms of life. Nature 582, 592–596 (2020).

19. Ronneberger, O., Fischer, P. & Brox, T. U-Net: Convolutional Networks for Biomedical Image Segmentation. ArXiv150504597 Cs (2015).

20. Kipf, T. N. & Welling, M. Semi-Supervised Classification with Graph Convolutional Networks. ArXiv160902907 Cs Stat (2017).

21. AlQuraishi, M. ProteinNet: a standardized data set for machine learning of protein structure. BMC Bioinformatics 20, 311 (2019).

22. Kryshtafovych, A., Schwede, T., Topf, M., Fidelis, K. & Moult, J. Critical assessment of methods of protein structure prediction (CASP)—Round XIII. Proteins Struct. Funct. Bioinforma. 87, 1011–1020 (2019).

23. Dunham, A., Jang, G. M., Muralidharan, M., Swaney, D. & Beltrao, P. A missense variant effect prediction and annotation resource for SARS-CoV-2. bioRxiv (2021) doi:10.1101/2021.02.24.432721.

24. Yamada, K. D. & Kinoshita, K. De novo profile generation based on sequence context specificity with the long short-term memory network. BMC Bioinformatics 19, 272 (2018).

25. Dunham, A. S. & Beltrao, P. Exploring amino acid functions in a deep mutational landscape. Mol. Syst. Biol. 17, e10305 (2021).

26. Jelier, R., Semple, J. I., Garcia-Verdugo, R. & Lehner, B. Predicting phenotypic variation in yeast from individual genome sequences. Nat. Genet. 43, 1270–1274 (2011).

27. Meier, J. et al. Language models enable zero-shot prediction of the effects of mutations on protein function. http://biorxiv.org/lookup/doi/10.1101/2021.07.09.450648 (2021).

28. The 1000 Genomes Project Consortium. A global reference for human genetic variation. Nature 526, 68–74 (2015).

29. Drummond, D. A., Bloom, J. D., Adami, C., Wilke, C. O. & Arnold, F. H. Why highly expressed proteins evolve slowly. Proc. Natl. Acad. Sci. U. S. A. 102, 14338–14343 (2005).

30. Wagih, O. et al. A resource of variant effect predictions of single nucleotide variants in model organisms. Mol. Syst. Biol. 14, e8430 (2018).

31. Liu, X., Jian, X. & Boerwinkle, E. dbNSFP: a lightweight database of human nonsynonymous SNPs and their functional predictions. Hum. Mutat. 32, 894–899 (2011).

32. Liu, X., Li, C., Mou, C., Dong, Y. & Tu, Y. dbNSFP v4: a comprehensive database of transcript-specific functional predictions and annotations for human nonsynonymous and splice-site SNVs. Genome Med. 12, 103 (2020).

33. Senior, A. W. et al. Improved protein structure prediction using potentials from deep learning. Nature 577, 706–710 (2020).

34. Shen, Z., Bao, W. & Huang, D.-S. Recurrent Neural Network for Predicting Transcription Factor Binding Sites. Sci. Rep. 8, (2018).

35. Pan, X., Rijnbeek, P., Yan, J. & Shen, H.-B. Prediction of RNA-protein sequence and structure binding preferences using deep convolutional and recurrent neural networks. BMC Genomics 19, 511 (2018).

36. Jing, B., Eismann, S., Soni, P. N. & Dror, R. O. Equivariant Graph Neural Networks for 3D Macromolecular Structure. ArXiv210603843 Cs Q-Bio (2021).

37. Fout, A., Byrd, J., Shariat, B. & Ben-Hur, A. Protein Interface Prediction using Graph Convolutional Networks. in 10 (2017).

38. Zamora-Resendiz, R. & Crivelli, S. Structural Learning of Proteins Using Graph Convolutional Neural Networks. bioRxiv 610444 (2019) doi:10.1101/610444.

39. Brown, T. B. et al. Language Models are Few-Shot Learners. ArXiv200514165 Cs (2020).

40. The Darwin Tree of Life Project Consortium. Sequence locally, think globally: The Darwin Tree of Life Project. Proc. Natl. Acad. Sci. 119, e2115642118 (2022).

41. Grealey, J. et al. The Carbon Footprint of Bioinformatics. Mol. Biol. Evol. 39, msac034 (2022).

