## Supplementary Information for "High-throughput deep learning variant effect prediction with Sequence UNET"

### UNET Model

The Sequence UNET model is implemented in TensorFlow[1] (version 2.5+) using the Keras framework. It takes a 3-dimensional tensor ($N \times M \times 20$) representing one-hot encoded protein sequences as input, with the final dimension being the amino acid, the middle protein position and the first outer dimension batches. As shown in **Fig. 1**, this is fed through $L$ compression CNN layers, where each layer contains two 1D CNN operations with $f \times 2^l$ filters on layer $l$ and width $k$ kernels. 1D max pooling operations compress the output of higher layers in half to input into the next layer. The input must be 0-padded in the second dimension, so the length is divisible by 2 $L-1$ times. The bottom layer contains a third CNN operation. $L$ corresponding expansion layers form the opposite side of the U structure, which each take input from a concatenation of the output from the corresponding compression layer and up-sampled output from the preceding expansion layer or the bottom layer. These also include 2 1D CNN operations. A final classification head processes the output of the final expansion layer, using an appropriate activation function for each task. Frequency classification uses a sigmoid function and frequency prediction the softmax function over the frequencies at each position. The final hyperparameters were $L = 6, f = 64, k = 9$, with ReLu activation functions for the CNN layers. The Swish activation function[2] was found to have slightly higher performance but technical details of TensorFlow 2.5 meant model gradients could not be loaded using Swish. In the future it could slightly improve performance but currently ReLU has been used to make distribution easier. Dropout at frequency 0.05 and batch normalisation were used in between each compression/expansion layer.

#### Structural Input

An optional simple GraphCNN[3] (**Eqn. (1)**) is available to encode per position protein structure information, which is then concatenated with the one hot encoded sequence and fed into the main model. In this case a weighted residue distance matrix (**Eqn. (2)**) is required as an additional input. We tested various sizes, number of graph layers and residue distances weightings but found a single graph layer with 32 filters and ELu activation performed best.

$$H^{i+1} = \sigma\left(AH^iW^i\right) \tag{1}$$

$A$ is the $n \times n$ normalised edge matrix, which must be normalised to prevent the overall magnitude of features from changing and causing exploding or vanishing gradients. $H^i$ is an $n \times m$ matrix of the hidden values in layer i, for a graph with n nodes with m features. $W^i$ is an $m \times l$ matrix of learnt weights for layer i, where $l$ is the number of features calculated for the next layer. This gives positional output that is a weighted sum of features from that and neighbouring amino acids, and operates independently of the input protein size.

The contact graph considers all residues within 10Å to be in contact, and uses a similarity metric (**Eqn. (3)**) to weight closer residues more highly. A self-connection was used to pass information about the position itself and residues one or two positions away in sequence that were missing structural information were assumed to be 380nm or 610nm away, based on the average of non-masked positions. Other masked residues were assumed to not be in contact. This was one of the less explored areas of the network, so it is likely more complex structural features could improve performance.

$$a_{ij} = \frac{c_{ij}}{\sum_k c_{ik}} \tag{2}$$

$$c_{ij} = \begin{cases} \dfrac{1}{3 + d_{ij}}, & d_{ij} \leq 10 \\ 0, & d_{ij} > 10 \end{cases} \tag{3}$$

where $d_{ij}$ is the distance between residues $i$ and $j$, $c_{ij}$ is the closeness between residues and $a_{ij}$ is the normalised residue contact matrix used in **Eqn. (1)**. Distances are measured in angstroms. The additional 3Å means self-connections are weighted as roughly twice that of the nearest neighbour.

### ProteinNet Dataset

We train the model using the ProteinNet dataset[4], which contains training, validation and test sets based on the CASP competition[5], with test sets containing the proteins predicted for each round of CASP and training sets containing all structures available in the PDB at the time of each competition. This does potentially introduce a slight bias into the test sets since only proteins with newly discovered crystal structures are included in CASP competitions, meaning they tend to be more unusual and less well studied proteins or those that are difficult to crystallise. However, the test sets still contain diverse proteins from a range of organisms, so any reduction in generalisation is likely to be small and predictions would generally be expected to be worse on these unusual proteins that cover novel sequence space. We use the CASP12 95% thinned training set as a balance between including a large variety of slightly different sequences and moderating the dataset size for rapid training. CASP is designed to assess template-based structure prediction methods[6] so a minority of proteins in the test set (38 of 146 in CASP12) have some sequence similarity to proteins in the PDB, which could potentially artificially inflate test performance by a small amount. However, the size of the training dataset means the model is unable to memorise variant consequences and so having a minority of somewhat related sequences in the test set is very unlikely to significantly influence results. Together, this makes ProteinNet a convenient and appropriate dataset for predicting deleteriousness from sequence and structure, despite that not being its original purpose, and using it greatly sped up model development.

### Training and Hyperparameter Optimisation

The model was trained and assessed using 32GB Nvidia Tesla V100 GPU nodes on a high-performance computer cluster. ProteinNetPy was used to generate the Tensorflow datasets for training and validation, using custom map functions depending on the model configuration being trained. A progressive hyperparameter optimisation strategy was used, testing a range of values for each parameter and using the best results as the default value of that parameter for future tests. The tested parameters were number of UNET layers, number of filters, kernel width, presence of structure features, optimisation algorithm, deleteriousness threshold, activation function and regularisation regime (**Fig. S1**). These were tested for both PSSM prediction and frequency classification modes of the model, with the same resulting parameters. When not testing optimisers or their settings, training was performed using the Adam optimiser[7] with a learning rate of 0.01 and early stopping based on validation accuracy with a memory of 20 epochs. Increasing model size tends to increase performance, so training speed and resource consumption is balanced directly against performance. We settled on 64 first layer filters, doubling in each layer; 9 wide kernels; and 6 layers to comfortably train on a single Nvidia Tesla V100 GPU; the size could be increased using more resources to incrementally improve performance. Increasing the number or size of GraphCNN layers did not improve performance, suggesting more advanced techniques[8,9] might be required to fully harness structure. A single 32 filter GraphCNN layer was used to encode structural features.

Three sizes of baseline convolutional model were also trained for comparison: a single layer network with 32 7 width filters; an equivalent double layer network; and a larger network with 64 7 width filters in the second layer. Training was performed using the CASP12 section of the ProteinNet dataset, using the 95% data split. The training code available at github.com/allydunham/sequence_unet contains the exact specification of the training procedures used.

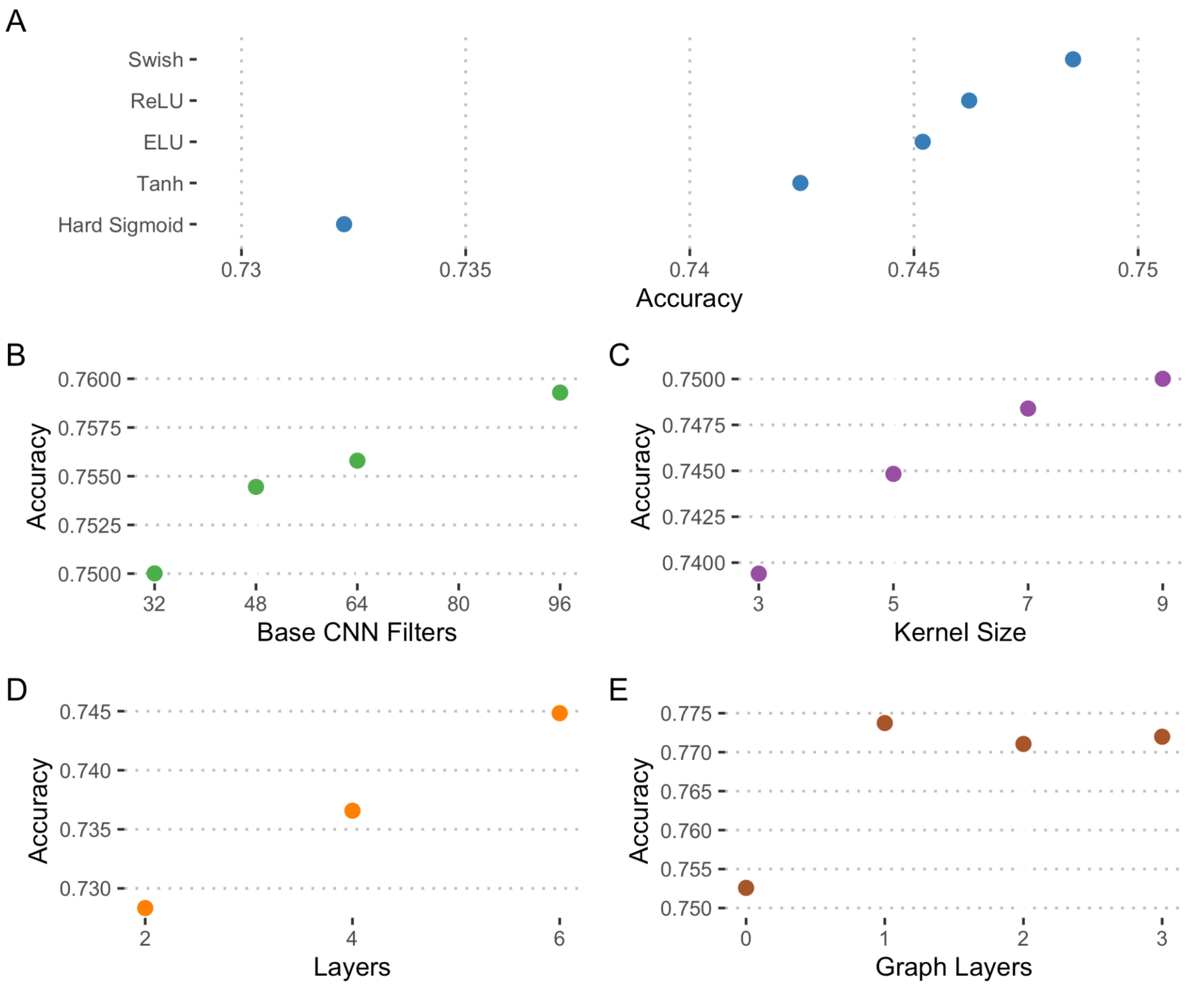


**Figure S1** - Hyperparameter optimisation. These results are from optimising the frequency classification model, but equivalent results occurred when optimising the PSSM predictor. **A**: CNN activation function. **B**: Number of CNN filters. **C**: Size of CNN kernel. **D**: Number of UNET compression layers. **E**: Number of graph CNN layers.

### Position specific variant frequency prediction

We first trained Sequence UNET to predict the frequency of each possible mutation in a sequence, and therefore its PSSM. This matrix compares the prevalence of each amino acid at each position to background amino acid frequencies, giving a measure of how well each substitution would be tolerated. The Kullbeck-Liebler divergence between the vector of predicted and true frequencies of each position was used as the loss function and a softmax activation function was used in the final layer to ensure the variant frequencies at each position sum to 1. The PSSMs predicted by this model are very similar to those derived from multiple sequence alignments (**Fig. S2A-B**). The average difference between predicted and true values in the ProteinNet CASP12 test set was 0.04, although this is quite variable ($\sigma = 0.072$). Interestingly, it is often the wild-type amino acid that is most incorrectly predicted (WT: $\mu = 0.18$, $\sigma = 0.184$, Missense: $\mu = 0.033$, $\sigma = 0.05$), which suggests the model is not able to learn how common the wild-type is as well as it learns what variants are tolerated or rejected.

SPBuild[10] is a recurrent neural network model that makes PSSM predictions and was shown to outperform similar de novo profile generation methods such as CSBuild[11,12] and RPS-Blast[13], so provides a good comparison for state of the art PSSM prediction from sequence. It is also a good general comparison between Sequence UNETs architecture and recurrent models, which were previously the standard approach to sequence-based problems. We generated SPbuild predictions for all proteins in the ProteinNet CASP 12 test set using the latest version of SPBuild (2020-01-07).

We also compare to ESM-1b[14,15], which provides a good comparison to large transformer based general protein language models. These are trained by learning to predict which amino acid occurs at each position, meaning they produce a vector of propensities for each amino acid at each position that relates to the PSSM. However, language models are generally used for the position representations they produce, which are used as a general base for task specific models. To compare to this usage we also trained a simple softmax layer to predict PSSM frequencies from ESM-1b representations, based on the ProteinNet CASP12 95% thinning data.

Sequence UNET naturally outputs raw variant frequencies, whereas PSSMs are generally reported as log scores normalised against wild-type amino acid frequencies (**Eqn. (4)**), which is what SPBuild reports. The frequencies output by Sequence UNET, the baseline CNN and ESM-1b models were transformed into the standard PSSM format to compare them to BLOSUM62 and SPBuild. Sequence UNET and the baseline CNN results were generated from the ProteinNet datafile using the sequence_unet package.

$$\log_2 \frac{f_{pred} + 10^{-5}}{f_{AA}} \quad (4)$$

Where $f_{pred}$ is the predicted frequency and $f_{AA}$ the average frequency of that amino acid, based on Swiss-Prot summary statistics[16,17].

The correlation between predictions and true values in the ProteinNet test set (**Fig. S2C**) suggests Sequence UNET ($\rho = 0.451$) outperforms SPBuild ($\rho = 0.332$), baseline evolutionary expectations ($\rho = 0.333$) and ESM-1b (top model $\rho = 0.237$, logits $\rho = 0.057$), with structural features increasing correlation slightly ($\rho = 0.472$). Interestingly base ESM-1b logits strongly correlate with variant frequencies ($\rho = 0.551$) but not the PSSM scores, which are normalised against overall amino acid frequencies. This may be because it is trained to identify the most likely amino acid and cannot differentiate magnitudes of rare variants, which are more important for PSSM scores. Sequence UNET and the ESM-1b top model are consistently most likely to be within 1 unit of the true value, be closest to the true value or both (**Fig. S2D**). ESM-1b performs better on these metrics than Pearson correlation, perhaps because it often correctly predicts the frequent low magnitude PSSM scores but fails to predict the high impact variants, which have more impact on correlation than these metrics. These results demonstrate that Sequence UNET improves on the current state of the art for specific de novo PSSM profile prediction and show that the model architecture can outperform LSTM recurrent models while being much more computationally efficient. They also suggest the model compares favourably to much more computationally intensive language models on this task.

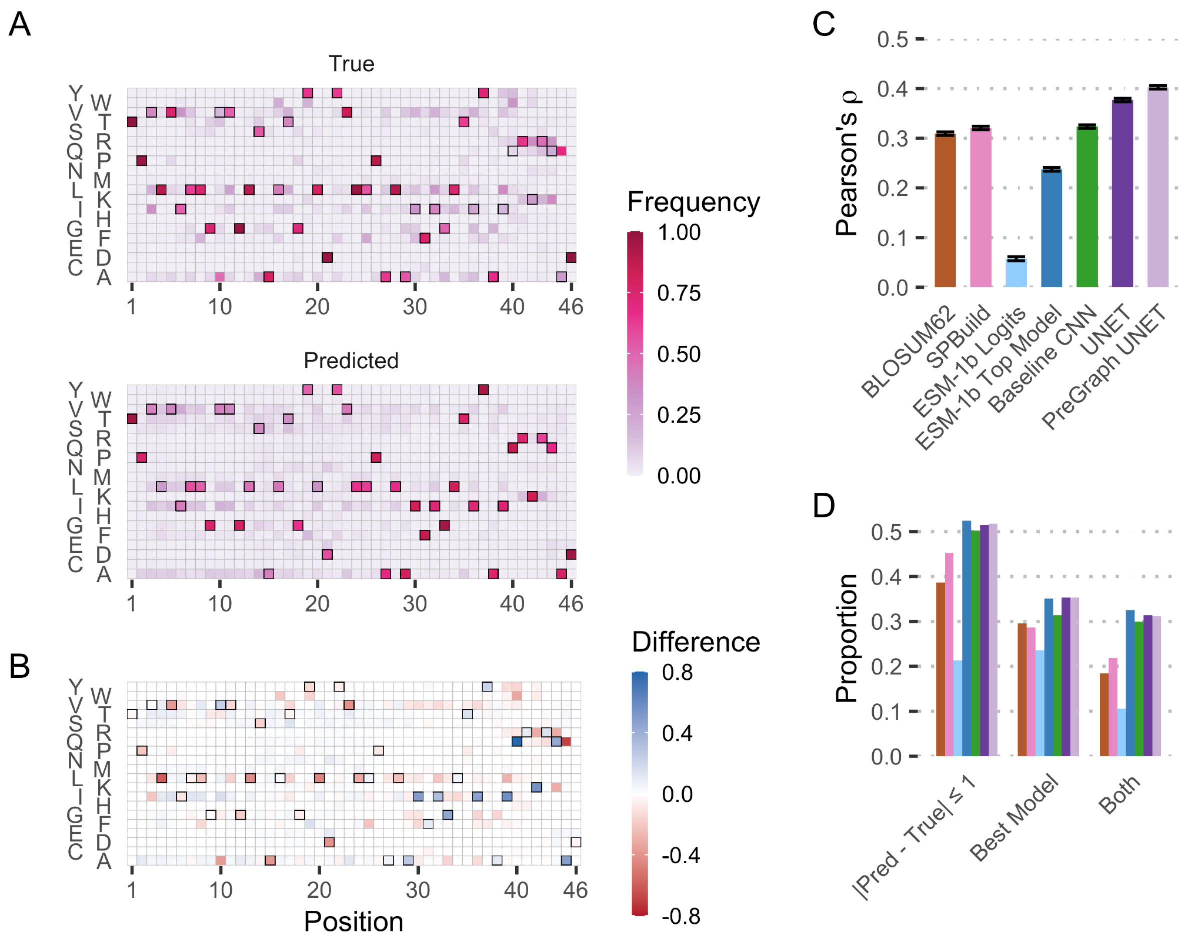


**Figure S2** - PSSM Prediction **A**: Example true and predicted MSA frequency predictions from the Sequence UNET model for the bacterial mercury transporter MerF. The wild-type amino acid at each position is outlined. **B**: Difference between predicted and true values of the PSSMs in **A**. **C**: Pearson correlation between predicted and true PSSM values for Sequence UNET, a single layer CNN, SPBuild, ESM-1b logits, an ESM-1b top model and BLOSUM62 based on the CASP12 test set. Confidence intervals are based on Fisher's Z transform of the correlation coefficient. **D**: PSSM prediction comparison showing proportion of predictions within 1 unit of the true value, that are the best or equal best and both. The models are coloured as in **C**.

### Variant frequency classification

The model can also be trained as a frequency classifier, using a binary cross-entropy loss function and a sigmoid activation function in the final layer. This outputs a score predicting whether each variant occurs below a specific frequency threshold (**Fig. S3A**), which can be treated as a soft deleteriousness score under the assumption that deleterious variants occur at low frequencies across homologous proteins. There is a significant separation between high and low frequency variants in test set proteins (**Fig. S3B**).

We compared classification performance with: a baseline single layer CNN; BLOSUM62 scores to represent an evolutionary baseline; SIFT4G[18] scores, which are based on an MSA and approximate deleteriousness; ESM-1b logits; and a single layer ESM-1b representation top model trained on CASP12 95% thinned ProteinNet data. We only compare to one VEP (SIFT4G) because while variant frequency is related to deleteriousness it is not exactly the same, meaning most VEPs are not directly comparable. We compare the generalised pathogenicity predictor model more widely. SIFT4G results were generated for the ProteinNet test set using default parameters and aligning against UniRef90. Sequence UNET and baseline CNN results were again generated using the python package. The single sigmoid layer ESM-1b top model was again trained on ProteinNet CASP12 95% thinned training data and scores predicted from the test set as well as ESM-1b logit results.

Sequence UNET matches the ESM-1b top model and outperforms all other models on the ProteinNet test set proteins when using models' natural deleteriousness thresholds. Again GraphCNN features slightly improve performance (**Fig. S3C**). It has higher average accuracy and precision than the other predictors as well as a higher median F1 score, indicating a better balance between precision and recall. This analysis highlights that SIFT4G is tuned for recall, having very high recall but low precision. Finally, Sequence UNET has a higher average Cohen's κ score, indicating performance that is further improved over random chance. In contrast, BLOSUM62, SIFT4G and ESM-1b logits can be little better than chance for many proteins.

Different frequency thresholds lead to very different training outcomes (**Fig. S3D**), with a difference in accuracy over 10% between tested thresholds. The difference in performance likely reflects the difficulty of determining the status of intermediate frequency variants, which are not obviously incompatible amino acids but also do not follow common motifs. Different thresholds can be potentially valuable for different applications, for example the 0.1 threshold could be useful for identifying variants very likely to be neutral and conversely low thresholds can identify very rare mutations that are likely to be deleterious. We primarily used a threshold of < 0.01 for further tests, reasoning that this is a good cut-off for general deleteriousness and that optimisations to this problem, which appears to be the most difficult, will carry over to other thresholds as well.

A ROC analysis gives a better overview of overall predictor characteristics, testing performance over a range of deleteriousness thresholds. The final model has a ROC AUC of 0.81 (**Fig. S3E**), making it much better at frequency classification than SIFT4G and BLOSUM62 scores (both AUC = 0.61). Adding structural features again improves performance a small amount (AUC = 0.83), resulting in a very similar performance profile to the ESM-1b top model (AUC = 0.83). A similar pattern is observed in precision recall analysis, with our model performing best (auPR = 0.79/0.81) alongside the ESM-1b top model (auPR = 0.8) (**Fig. S3F**). SIFT4G likely performs poorly on this task, compared with reasonable performance elsewhere, because the SIFT4G score is calculated by normalising modelled MSA frequency against the frequency of the most frequent amino acid at that position, which allows a convenient consistent deleteriousness cut-off across positions but throws away information about the absolute value of frequencies. Scores from EVE and DeepSequence would also likely suffer from this problem. Conversely ESM-1b logits seem likely to be good at determining likely amino acids at a position but are poor at determining those that are particularly rare, explaining poor PSSM and low frequency classification performance.

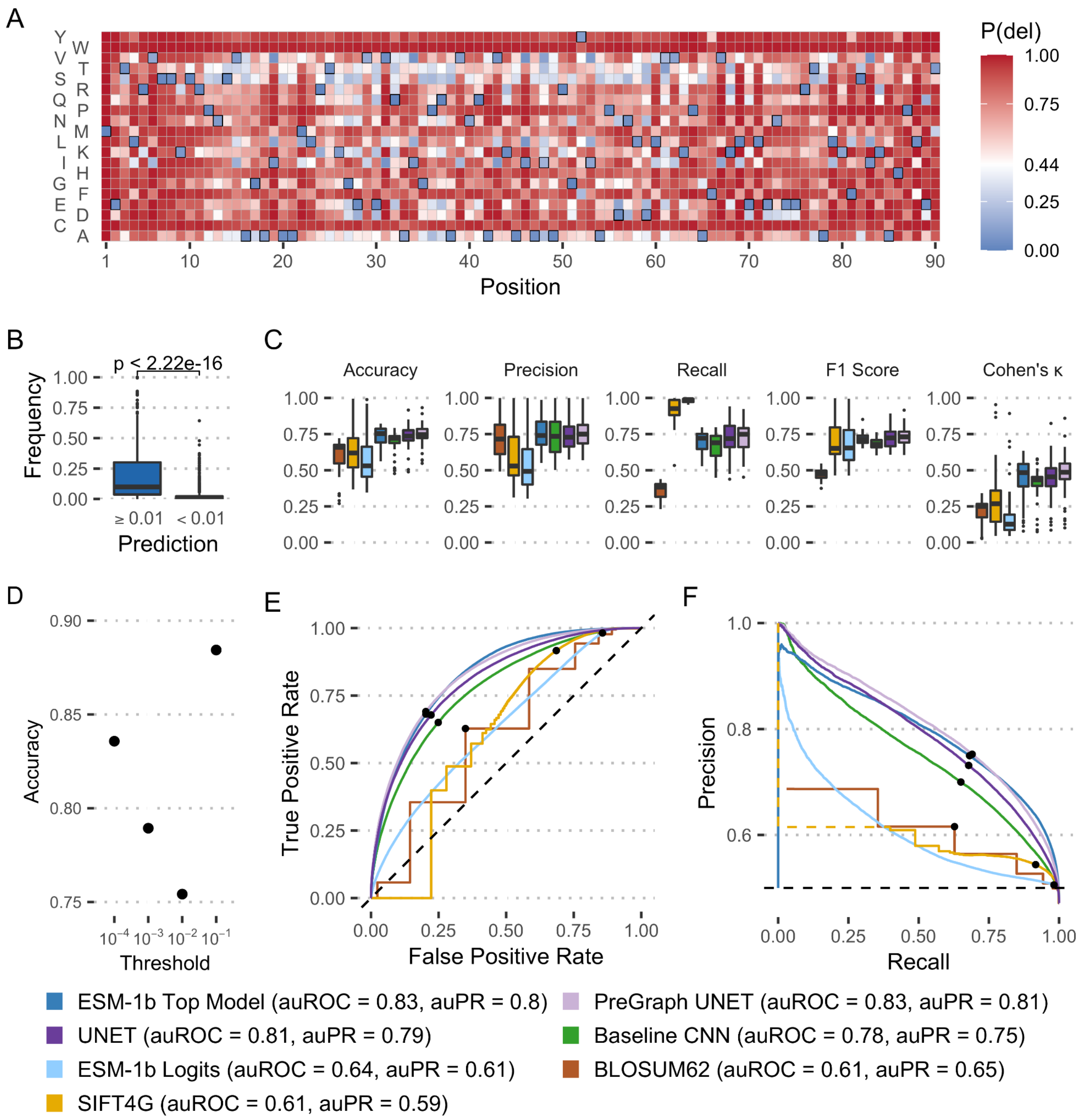


**Figure S3** - Variant Frequency Classification **A**: Example of sequence-based frequency classification (f < 0.01) on *T. thermophilus* protein SPOVS. The deleteriousness threshold (P(del) > 0.44) was chosen to balance specificity and sensitivity by maximising $\sqrt{(1-FPR)^2+TPR^2}$(see **E**). **B**: Observed MSA frequency distribution of variants in SPOVS predicted to be above and below 0.01 frequency. **C**: Performance statistics comparing Sequence UNET with and without structural features, ESM-1b logits, an ESM-1b top model, a baseline single layer CNN, SIFT4G and BLOSUM62. The first four are at their natural deleteriousness thresholds (0.5 for the neural networks & 0.05 for SIFT4G) and the optimal threshold for BLOSUM62 was determined to be -2 based on the ROC analysis. Each boxplot shows the distribution of performance statistics over the different proteins in the ProteinNet Casp12 95% thinned test dataset. **D**: Frequency classification accuracy of models trained at different deleterious frequency thresholds. **E/F**: ROC (**E**) and PR (**F**) curves comparing Frequency classification (< 0.01) performance of the models in **C**. The standard model deleteriousness thresholds are marked with black points on the curves.

### Model Generalisation

Generalisation was tested using ClinVar[19] (24/04/2021 dataset), a standardised DMS dataset we previously generated[20] and a set of gold standard *S. cerevisiae* variants[21]. ClinVar classifications were simplified to pathogenic or benign, each category including variants designated as likely or definitely benign/pathogenic. DMS ER scores less than -0.5 were classified as deleterious. The standard models were tested against these datasets and fine-tuned models were trained to predict ClinVar pathogenicity, either by replacing the classification head layer or refining its weights with the rest of the network frozen. Fine-tuned models in both styles were trained based on both PSSM prediction and frequency classification with 3 and 1 width convolution kernels. They were trained using binary cross entropy loss and the Adam optimiser for a maximum of 50 epochs with 10 epoch memory early stopping. In practice validation performance decreased after 10 to 15 epochs in each case, so weights from this stage were selected by the early stopping procedure. Three wide kernel models based on the classification model performed best and were used.

The ClinVar dataset has a narrow focus, containing information on human variants with validated clinical significance, leading to biases in protein composition with less than 2000 well studied human proteins represented and 14 proteins each with over 100 variants together accounting for 15.4% of the dataset variants. Despite these biases a random split was sufficient to get variants from a broad range of proteins in each set, with proteins and positions exclusively in each set and no single protein dominating. This suggests the model can not just learn the properties of individual positions or proteins, especially as it is only trained for a few epochs. However, the limited range and type of protein included in the dataset does suggest that the fine-tuning will not necessarily transfer to other families of proteins or those from other species.

SIFT4G and FoldX[22] predictions for ClinVar variants were retrieved from Mutfunc[23], ESM-1v scores were calculated directly, EVE scores were downloaded from its website and all other tool results were retrieved from dbNSFP[24,25].

### Prediction Variation with Variable Padding

The model supports padding sequences with additional zeroes to allow proteins of different lengths to be processed in batches; a fact that is utilised to speed up training and can speed up predictions. This is in addition to the mandatory padding required to make most proteins lengths divisible by $2^5$. Additional padding does alter prediction calculation, but was included during training so the network would be expected to learn to account for variable length. This is generally found to be the case, although in a small number of cases changing padding changes variant predictions significantly. We tested the impact of padding by making predictions for proteins in the ProteinNet CASP12 validation set with 32 to 1024 amino acids of additional padding, testing both the frequency classifier and PSSM predictor models. In general, the padded predictions correlated very strongly with unpadded predictions (**Fig. S4A**). Correlation decreased a reasonable amount in a few shorter proteins, particularly for the classifier model, but predictions in the vast majority of proteins correlated with $\rho > 0.9$. Correlation also decreases, albeit modestly in most cases, as the amount of additional padding increases, suggesting it is beneficial to group similar length variants in batches to minimise the required padding. It is also rare for predictions to change classification, with the altered prediction crossing the 0.5 boundary (**Fig. S4B**). Again, it is shorter proteins that tend to have the most variants changing class. Overall, the average difference between padded and unpadded predictions is generally modest, particularly for the PSSM predictor (**Fig. S4C**). The majority of predictions (> 70%) from the classifier still change by less than 0.05 with over 85% less than 0.1. Finally, additional padding is found to have a negligible effect on overall accuracy (**Fig. S4D**), including at positions near to the end of the protein which are generally affected more by padding changes. Overall, adding additional padding to make batch predictions does have a large impact on a small number of predictions but is unlikely to impact overall results, especially in the large-scale analysis this tool is most suited to. This allowed us to use batching for our large-scale analyses without impacting predictions too much. In future, adapting training to batch together similar length proteins and using a similar strategy during prediction would help to reduce this effect even further. It might also be beneficial to add a minimum padding to standardise results on shorter proteins, which are generally most affected.

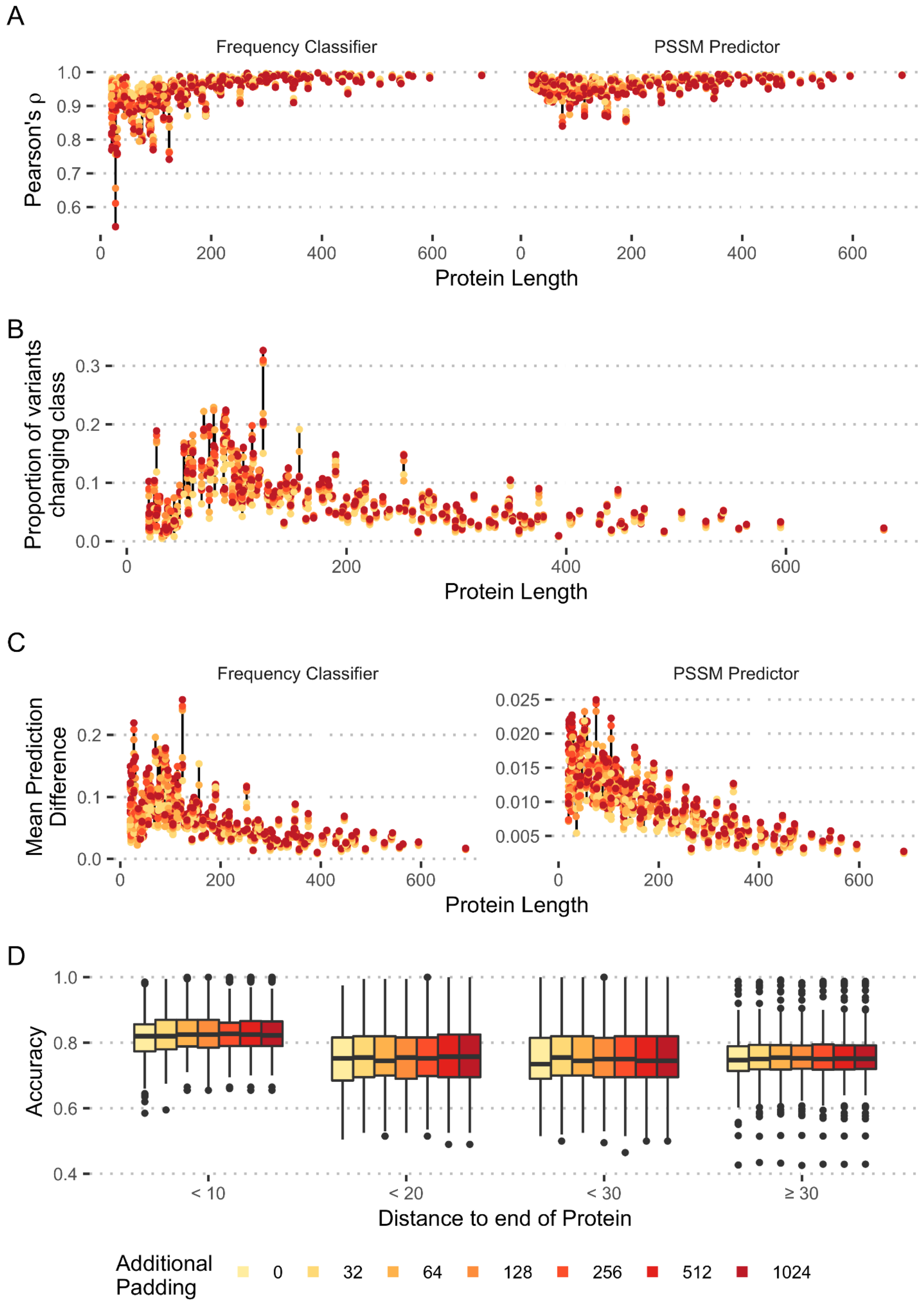


**Figure S4** - Impact of padding on predictions. **A**: Pearson correlation between unpadded predictions and those made on sequences with varying levels of additional padding. Correlations were calculated for each protein independently using both the PSSM predictor and frequency classifier model. **B**: Proportion of variants predicted to change class ($p \leq 0.5$

to $p > 0.5$ or vice-versa) when different levels of additional padding are added to sequences, using the frequency classifier model. **C**: Mean difference between unpadded and padded predictions on a range of proteins using both models. **D**: Accuracy of frequency classification in different proteins with various levels of additional padding, stratified by the distance between the position and the protein end, where padding begins. All panels are based on proteins from the ProteinNet CASP12 validation set. In **A**, **B** and **C** points from the same protein at different pad levels are linked by lines.

### Proteomic Analysis

The sequences of proteins identified in the Muller et al. pan-proteome analysis were downloaded from UniProt. The sequence_unet predict_from_fasta command was then used to generate Sequence UNET predictions using only sequence input for the PSSM prediction and frequency classification models. Prediction results for each protein were summarised as the mean number of predicted deleterious variants across positions. Protein abundance was normalised against length and expressed as the $\log_2$ fold change compared to median abundance in that species. The phylogeny of these organisms was downloaded from NCBI taxonomy[26,27]. SIFT4G scores for comparison were downloaded from Mutfunc[23] for *H. sapiens, S. cerevisiae* and *E. coli* and calculated using Uniref90 and default settings for Mycoplasma.

### References


1. Abadi, M. *et al.* TensorFlow: Large-Scale Machine Learning on Heterogeneous Distributed Systems. 19 (2015).
2. Ramachandran, P., Zoph, B. & Le, Q. V. Searching for Activation Functions. *ArXiv171005941 Cs* (2017).
3. Scarselli, F., Gori, M., Tsoi, A. C., Hagenbuchner, M. & Monfardini, G. The Graph Neural Network Model. *IEEE Trans. Neural Netw.* **20**, 61–80 (2009).
4. AlQuraishi, M. ProteinNet: a standardized data set for machine learning of protein structure. *BMC Bioinformatics* **20**, 311 (2019).
5. Kryshtafovych, A., Schwede, T., Topf, M., Fidelis, K. & Moult, J. Critical assessment of methods of protein structure prediction (CASP)—Round XIII. *Proteins Struct. Funct. Bioinforma.* **87**, 1011–1020 (2019).
6. Kryshtafovych, A. *et al.* Evaluation of the template-based modeling in CASP12. *Proteins* **86**, 321–334 (2018).
7. Kingma, D. P. & Ba, J. Adam: A Method for Stochastic Optimization. *ArXiv14126980 Cs* (2017).
8. Jing, B., Eismann, S., Soni, P. N. & Dror, R. O. Equivariant Graph Neural Networks for 3D Macromolecular Structure. *ArXiv210603843 Cs Q-Bio* (2021).
9. Jumper, J. *et al.* Highly accurate protein structure prediction with AlphaFold. *Nature* 1–11 (2021) doi:10.1038/s41586-021-03819-2.
10. Yamada, K. D. & Kinoshita, K. De novo profile generation based on sequence context specificity with the long short-term memory network. *BMC Bioinformatics* **19**, 272 (2018).
11. Biegert, A. & Söding, J. Sequence context-specific profiles for homology searching. *Proc. Natl. Acad. Sci. U. S. A.* **106**, 3770–3775 (2009).
12. Angermüller, C., Biegert, A. & Söding, J. Discriminative modelling of context-specific amino acid substitution probabilities. *Bioinformatics* **28**, 3240–3247 (2012).

13. Boratyn, G. M. *et al.* Domain enhanced lookup time accelerated BLAST. *Biol. Direct* **7**, 12 (2012).

14. Rives, A. *et al.* Biological structure and function emerge from scaling unsupervised learning to 250 million protein sequences. *Proc. Natl. Acad. Sci.* **118**, e2016239118 (2021).

15. Meier, J. *et al. Language models enable zero-shot prediction of the effects of mutations on protein function*. http://biorxiv.org/lookup/doi/10.1101/2021.07.09.450648 (2021).

16. Bienert, S. *et al.* The SWISS-MODEL Repository—new features and functionality. *Nucleic Acids Res.* **45**, D313–D319 (2017).

17. Waterhouse, A. *et al.* SWISS-MODEL: homology modelling of protein structures and complexes. *Nucleic Acids Res.* **46**, W296–W303 (2018).

18. Vaser, R., Adusumalli, S., Leng, S. N., Sikic, M. & Ng, P. C. SIFT missense predictions for genomes. *Nat. Protoc.* **11**, 1–9 (2015).

19. Landrum, M. J. *et al.* ClinVar: improving access to variant interpretations and supporting evidence. *Nucleic Acids Res.* (2018).

20. Dunham, A. S. & Beltrao, P. Exploring amino acid functions in a deep mutational landscape. *Mol. Syst. Biol.* **17**, e10305 (2021).

21. Jelier, R., Semple, J. I., Garcia-Verdugo, R. & Lehner, B. Predicting phenotypic variation in yeast from individual genome sequences. *Nat. Genet.* **43**, 1270–1274 (2011).

22. Schymkowitz, J. *et al.* The FoldX web server: an online force field. *Nucleic Acids Res.* **33**, W382–W388 (2005).

23. Wagih, O. *et al.* A resource of variant effect predictions of single nucleotide variants in model organisms. *Mol. Syst. Biol.* **14**, e8430 (2018).

24. Liu, X., Jian, X. & Boerwinkle, E. dbNSFP: a lightweight database of human nonsynonymous SNPs and their functional predictions. *Hum. Mutat.* **32**, 894–899 (2011).

25. Liu, X., Li, C., Mou, C., Dong, Y. & Tu, Y. dbNSFP v4: a comprehensive database of transcript-specific functional predictions and annotations for human nonsynonymous and

splice-site SNVs. *Genome Med.* **12**, 103 (2020).

26. Sayers, E. W. *et al.* GenBank. *Nucleic Acids Res.* **47**, D94–D99 (2019).

27. Schoch, C. L. *et al.* NCBI Taxonomy: a comprehensive update on curation, resources and tools. *Database J. Biol. Databases Curation* **2020**, baaa062 (2020).
